# Cortical TRPV1 modulates sensory processing through TRPA1-dependent mechanisms

**DOI:** 10.64898/2026.08.24.746688

**Authors:** Leena Amrutha, Saba Gharaei, Kyogo Sakai, Ehsan Arabzadeh, Ehsan Kheradpezhouh

## Abstract

Transient receptor potential vanilloid 1 (TRPV1) is classically recognised as a peripheral ion channel involved in nociception, but its physiological role in cortical sensory processing remains poorly understood. Here, we investigated whether cortical TRPV1 modulates tactile perception and sensory representations in the vibrissal primary somatosensory cortex (vS1) of mice. TRPV1 activation enhanced perceptual sensitivity, particularly for near-threshold tactile stimuli. In vS1, this was accompanied by stimulus-evoked responses that enhanced stimulus selectivity and discriminability. At the cellular level, activating TRPV1 increased excitability of pyramidal neurons in a TRPA1-dependent manner. Within vS1, TRPV1 activation reduced positive noise correlations and improved stimulus decoding through TRPA1-dependent changes in population activity. Together, these findings identify a TRPV1-TRPA1 signalling mechanism that regulates cortical gain, sharpens sensory representations and enhances perceptual sensitivity. Our results extend the established role of TRPV1 beyond peripheral sensory transduction to central sensory computation.

## Introduction

Transient receptor potential vanilloid 1 (TRPV1) is best known as a peripheral sensory ion channel that detects salient environmental and physiological stimuli. As a non-selective cation channel with high calcium (Ca²⁺) permeability, it plays a critical role in pain, inflammation and thermal sensation ^1–3^. Beyond these well-studied peripheral functions, TRPV1 also acts as a multimodal channel that helps sense and regulate internal physiological states, including signals from visceral organs ^4,5^. TRPV1 is also expressed throughout the central nervous system, including the hippocampus, cerebellum, hindbrain, olfactory bulb, midbrain and the cortex ^6–8^ where information is integrated. More recently, TRPV1 expression has been identified on cortical neurons, where it has been shown to influence neuronal excitability and synaptic transmission ^9–11^. Although cortical TRPV1 has been implicated in pathological processes, including chronic pain, neuroinflammation, affective disorders and neurodegenerative diseases ^12–17^ its physiological function in cortical circuits remains largely unknown.

The presence of TRPV1 in the cortex raises a fundamental question: why is an ion channel specialised for sensory detection expressed in the cerebral cortex? Despite evidence that cortical TRPV1 modulates neuronal excitability ^10,11^ whether it shapes representations of sensory stimuli and consequently influences perception remains unknown. Here we directly examine the role of cortical TRPV1 in the vibrissal primary somatosensory cortex (vS1) of awake mice. We show that TRPV1 activation enhances tactile perceptual sensitivity, particularly for near-threshold stimuli, while amplifying sensory-evoked cortical responses and improving stimulus discriminability. TRPV1 activation also increases the excitability of pyramidal neurons and alters population dynamics, providing a potential cellular and circuit basis for its effects on sensory representations and perceptual performance. We further identify co-expression and functional interaction between TRPV1 and TRP Ankyrin 1 (TRPA1) in vS1. Together, these findings indicate that TRPA1 is required for TRPV1-driven enhancement of pyramidal neuron excitability and to modulate cortical sensory responses.

## Results

### TRPV1 enhances perceptual sensitivity and sensory evoked responses

TRPV1 is expressed and functionally active in cortical neurons ^9–11^, however its contribution to shaping cortical sensory responses has yet to be elucidated. We therefore investigated how modulating cortical TRPV1 activity influences sensory perception and neuronal encoding within the vibrissal primary somatosensory cortex (vS1) of awake mice. To address this, we trained mice to report whisker vibrations of varying amplitudes (0-80 *μ*m) by licking a reward spout in a head-fixed behavioural paradigm. TRPV1 activity was modulated through local drug perfusion via a chronically implanted cannula targeting vS1. Mice were tested under three conditions: artificial cerebrospinal fluid (aCSF; control), TRPV1 agonist capsaicin (CAP), and antagonist 6-iodonordihydrocapsaicin (6-CAP; **Figure 1a**). Simultaneously, we recorded population activity from layer 2/3 neurons using two-photon Ca^2+^ imaging in GCaMP7f-expressing mice.

**Figure 1.**
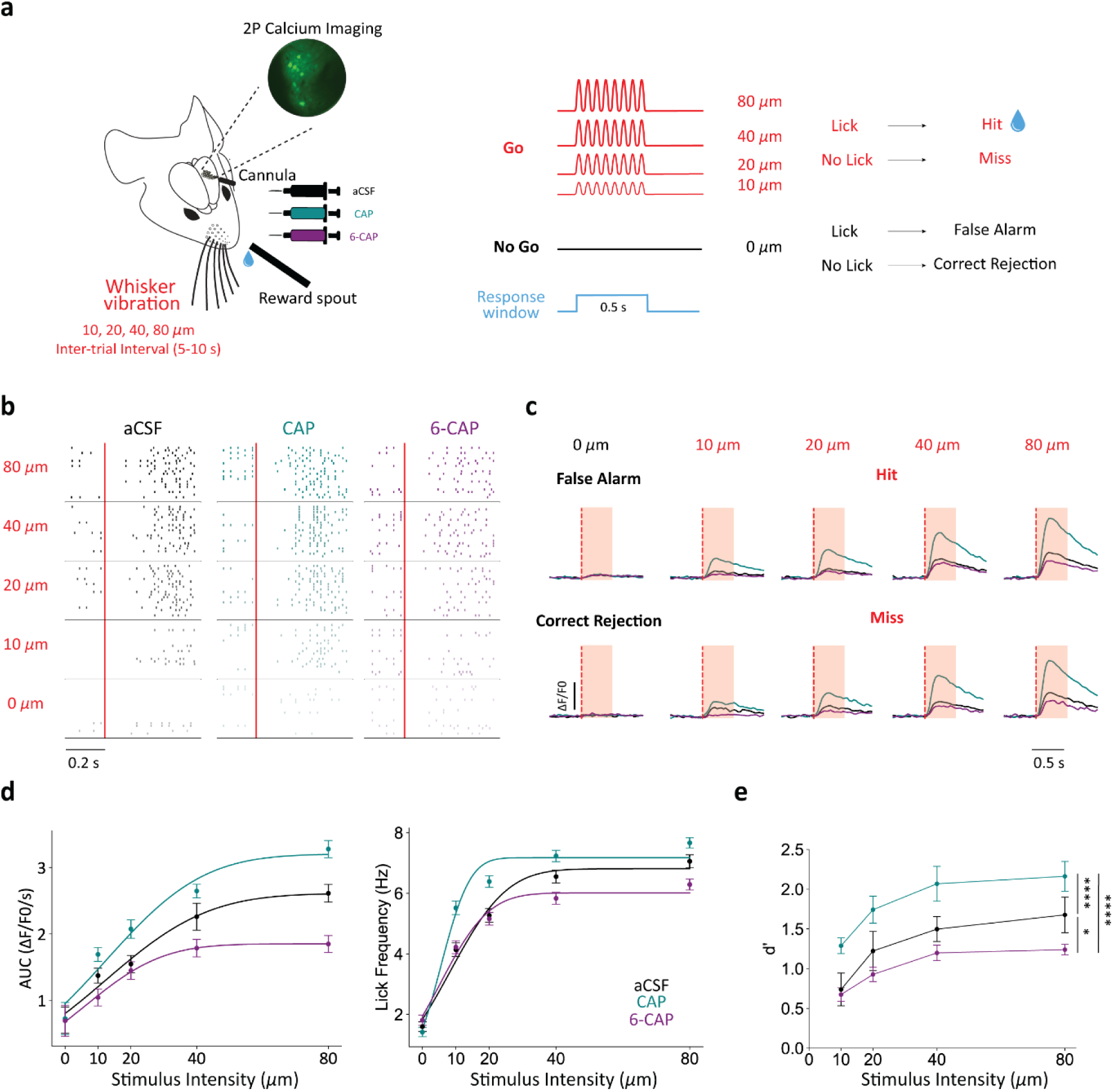
TRPV1 modulates sensory representations and perceptual sensitivity. **a)** Schematic of the head-fixed behavioural setup used for testing sensory perception. GCaMP7f-expressing mice received vibrational stimuli of varying amplitudes (0, 10, 20, 40, or 80μm) delivered to the left whisker pad via a piezoelectric mesh for 500ms. Mice reported stimulus detection by licking a reward spout to receive sucrose water (Hit trials). No lick during a stimulus trial was classified as a Miss; licking during a 0μm (catch) trial was a False Alarm (FA), and correctly withholding a lick was a Correct Rejection (CR). Neuronal activity in layer 2/3 of vibrissal primary somatosensory cortex (vS1) was recorded using two-photon Ca²^+^ imaging. Cortical TRPV1 activity was modulated via lateral perfusion of artificial cerebrospinal fluid (aCSF; Control), capsaicin (CAP; TRPV1 agonist), or 6-iodonordihydrocapsaicin (6-CAP; TRPV1 antagonist) through a chronically implanted cannula. **b)** Raster plots of lick responses from a representative mouse under aCSF (Control, black), CAP (teal), and 6-CAP (purple) conditions. Each dot indicates a lick, and each row represents an individual trial. Red vertical lines mark stimulus onset. **c)** Trial-averaged changes in Ca²⁺ fluorescence (ΔF/F) across all trial types and drug conditions (aCSF, CAP, and 6-CAP). Red dotted lines indicate stimulus onset. **d)** Neurometric curves (derived from the area under the curve [AUC] of Ca²^+^ responses within a 1s window post-stimulus) and corresponding psychometric curves (lick frequency) across all mice and drug conditions. **e)** Average d-prime (d′) across all mice for each stimulus intensity and condition. Data are presented as mean ± SEM. Statistical comparisons were performed using repeated-measures one-way ANOVA with Tukey’s post hoc test. *p < 0.05, ****p < 0.0001.

Modulation of TRPV1 produced systematic changes in detection behaviour. Under control conditions and consistent with established sensory-behavioural relationships, mice exhibited graded increases in lick probability as stimulus amplitude increased (**Figure 1b,d**). CAP induced a leftward shift in the psychometric function, indicating enhanced sensitivity to low-amplitude stimuli, whereas 6-CAP produced a rightward shift, consistent with diminished detection sensitivity. These effects were robust across animals and stimulus intensities (**Figure 1d *right***), and false alarm rates remained significantly unchanged across drug conditions (**Supplementary Figure 1b**). To quantify perceptual sensitivity independently of response bias, we computed d-prime (d′) across all stimulus amplitudes by comparing Hit and False Alarm rates. Mice consistently showed d’ values above 0 across all conditions. (**Supplementary Figure 1a**) confirming sustained and reliable task engagement. Notably, CAP produced a significant enhancement in d′ across the stimulus intensities, whereas 6-CAP reduced d’ values relative to control (**Figure 1e**; RM one-way ANOVA with Tukey’s post hoc, *p* < 0.0001 and *p* = 0.02, respectively), demonstrating that TRPV1 activity modulates perceptual sensitivity.

We next asked whether these behavioural effects were reflected in vS1 population activity. Neuronal responses were quantified as stimulus-evoked changes in fluorescence (ΔF/F), and the area under the curve (AUC, 0-1 s post-stimulus) was used to construct neurometric functions independently of behavioural outcome. Across all conditions, vS1 neurons exhibited amplitude-dependent increases in activity (**Figure 1c, d *left***). TRPV1 activation with CAP increased response magnitude across stimulus intensities, producing a leftward shift in the neurometric curve that mirrored behavioural improvements in detection. Conversely, TRPV1 inhibition attenuated sensory-evoked responses and produced a rightward shift in the neurometric function (**Figure 1d *left***). Baseline activity during 0 μm (catch) trials did not differ across drug conditions (n = 836 neurons, 6 mice, *p* > 0.05, Friedman test with Dunn’s post hoc), indicating that TRPV1 modulation primarily affected stimulus-evoked responses. Together, these results show a tight correspondence between perceptual sensitivity and sensory encoding in vS1 and are consistent with TRPV1-mediated gain-like modulation of cortical responses.

### TRPV1 activation enhances stimulus-selective encoding in vS1

We next examined whether TRPV1 modulation alters not only the magnitude but also the structure of sensory encoding in vS1. To distinguish stimulus-driven activity from activity related to behavioural choice, stimulus-balanced resampling showed that trial-to-trial variability in vS1 responses was positively associated with perceptual performance beyond the effects of stimulus intensity (**Supplementary Figure 2**). To characterise stimulus- and choice-related components of neuronal activity, we computed *stimulus index* and *choice index* for individual neurons across the three conditions (**Figure 2a**). These indices quantify the extent to which neuronal responses covary with the presence of the stimulus or with behavioural reports, respectively.

**Figure 2.**
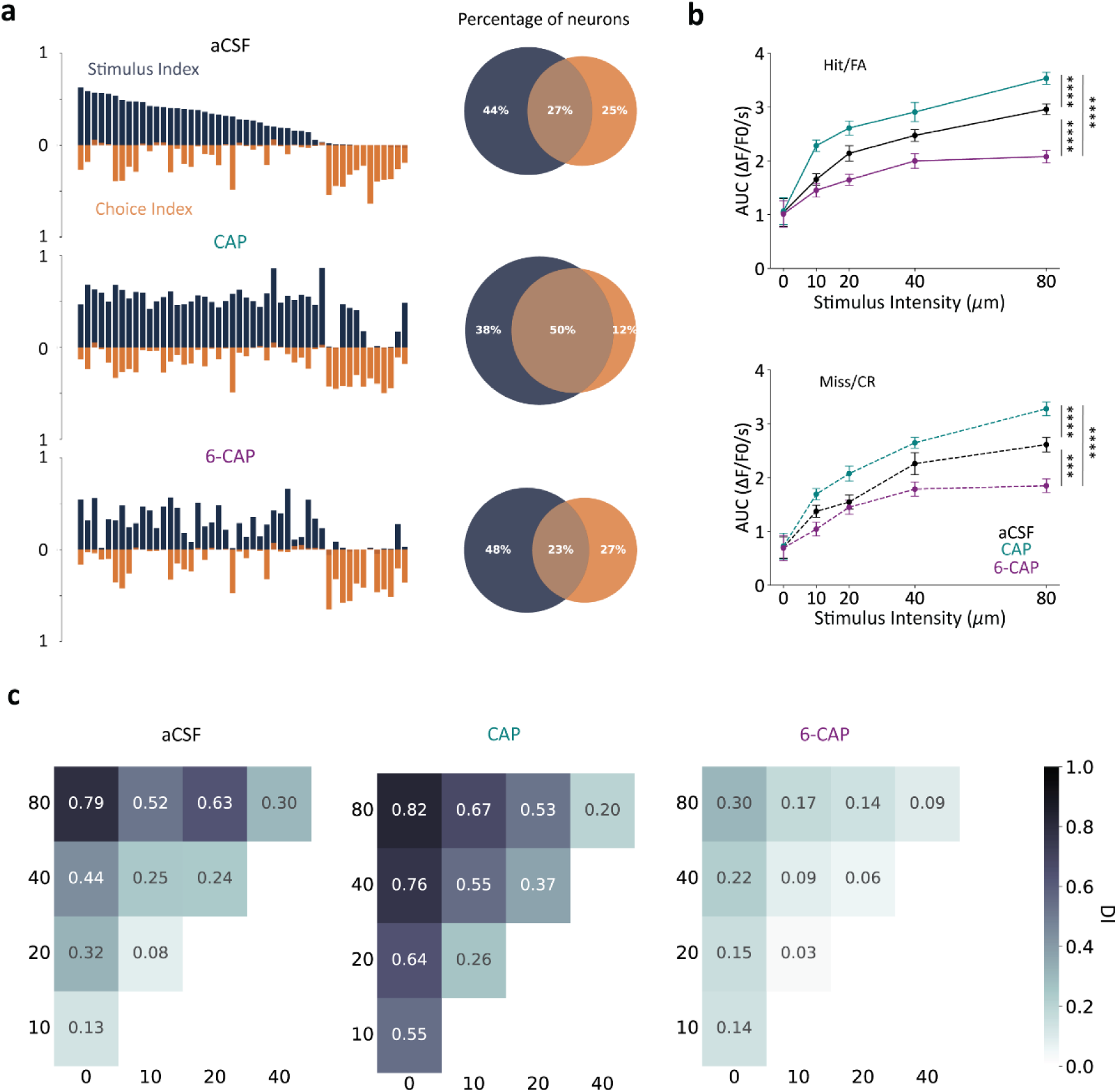
Stimulus and choice-related modulation by TRPV1 activation. **a)** Left: Bar plots showing the stimulus selectivity index (navy) and choice selectivity index (orange) for the same neurons across all drug conditions of one mouse. Neurons are sorted by stimulus selectivity under control conditions (aCSF) and tracked across capsaicin (CAP; TRPV1 agonist) and 6-CAP (TRPV1 antagonist) sessions. Right: Venn diagrams illustrating the proportion of neurons significantly modulated by stimulus (navy), choice (orange), or both, under each drug condition (n = 836 neurons from 6 mice). **b)** Top: Neurometric functions for Hit and False Alarm (FA) trials, measured by area under the curve (AUC) of Ca^2+^ responses (1 s post-stimulus), across all drug conditions. Bottom: Input–output functions for Miss and Correct Rejection (CR) trials, also measured by AUC across conditions. Data are presented as mean ± SEM. Statistical comparisons were performed using the Friedman test with Dunn’s multiple comparisons test. ***p < 0.001, ****p < 0.0001. **c)** Heatmaps of discrimination index (DI) values for individual neurons across all stimulus intensity pairs and drug conditions. Higher DI indicates greater neuronal discrimination between stimulus amplitudes.

Under control conditions (aCSF), most neurons (71%) were significantly modulated by stimulus amplitude, whereas a smaller subset (27%) exhibited activity correlated with behavioural choice. TRPV1 activation with CAP increased both the magnitude of stimulus selectivity and the fraction of stimulus-modulated neurons (88%), consistent with the enhanced sensory-evoked responses observed at the population level (**Figure 1c, d**). A subset of neurons previously classified as exhibiting choice-correlated activity under control conditions also showed stimulus selectivity under CAP. In contrast, TRPV1 inhibition with 6-CAP left the proportion of stimulus-responsive neurons largely unchanged relative to aCSF but reduced the magnitude of stimulus selectivity, in line with the observed attenuation of neurometric responses. Across all conditions, the proportion and strength of choice-correlated activity remained comparatively stable (**Figure 2a *right***), indicating that TRPV1 modulation predominantly affects stimulus encoding rather than choice-related activity.

To further determine whether TRPV1-dependent modulation of vS1 activity varied with behavioural outcome, we analysed neural responses separately for trials with and without lick responses. CAP enhanced stimulus-evoked activity across both lick (Hit/FA) and no-lick (Miss/CR) trials (n = 836 neurons, Friedman test with Dunn’s post hoc), whereas 6-CAP reduced responses in both categories (**Figure 2b**). These effects were present regardless of behavioural outcome, suggesting that the modulation was largely independent of licking and consistent with a predominantly stimulus-driven effect. To quantify representational precision while minimising behavioural confounds, we computed a *discrimination index* (DI) for all pairs of stimulus amplitudes using activity from Miss and Correct Rejection trials (**Figure 2c**). Under control conditions, DI values increased with stimulus separation, with stronger discrimination observed for more widely separated amplitudes. TRPV1 activation led to a marked enhancement in discriminability, particularly for adjacent low-amplitude stimulus pairs (10-20 *μ*m). This enhancement was less pronounced at higher amplitudes, consistent with response saturation observed in neurometric functions. In contrast, TRPV1 inhibition significantly reduced discriminability across most stimulus pairs. These results indicate that TRPV1 activity enhances the precision of stimulus encoding in vS1, particularly for weak tactile inputs.

### TRPV1-TRPA1 interactions modulate the excitability of cortical pyramidal neurons

Given the enhanced sensory responsiveness and perceptual sensitivity observed under TRPV1 activation, we next investigated the cellular mechanisms through which TRPV1 modulates cortical excitability in vS1. In peripheral sensory neurons, TRPV1 frequently co-expresses with TRPA1 and exhibits functional interactions through shared intracellular signalling pathways ^18,19^. We therefore asked whether a similar interaction exists in the cortex and whether TRPA1 contributes to TRPV1-dependent modulation of excitability. Immunohistochemical analysis revealed robust somatic co-expression of TRPV1 and TRPA1 in layer 2/3 pyramidal neurons of vS1 (**Figure 3a**; **Supplementary Figure 3**), providing anatomical support for potential intracellular interactions.

**Figure 3.**
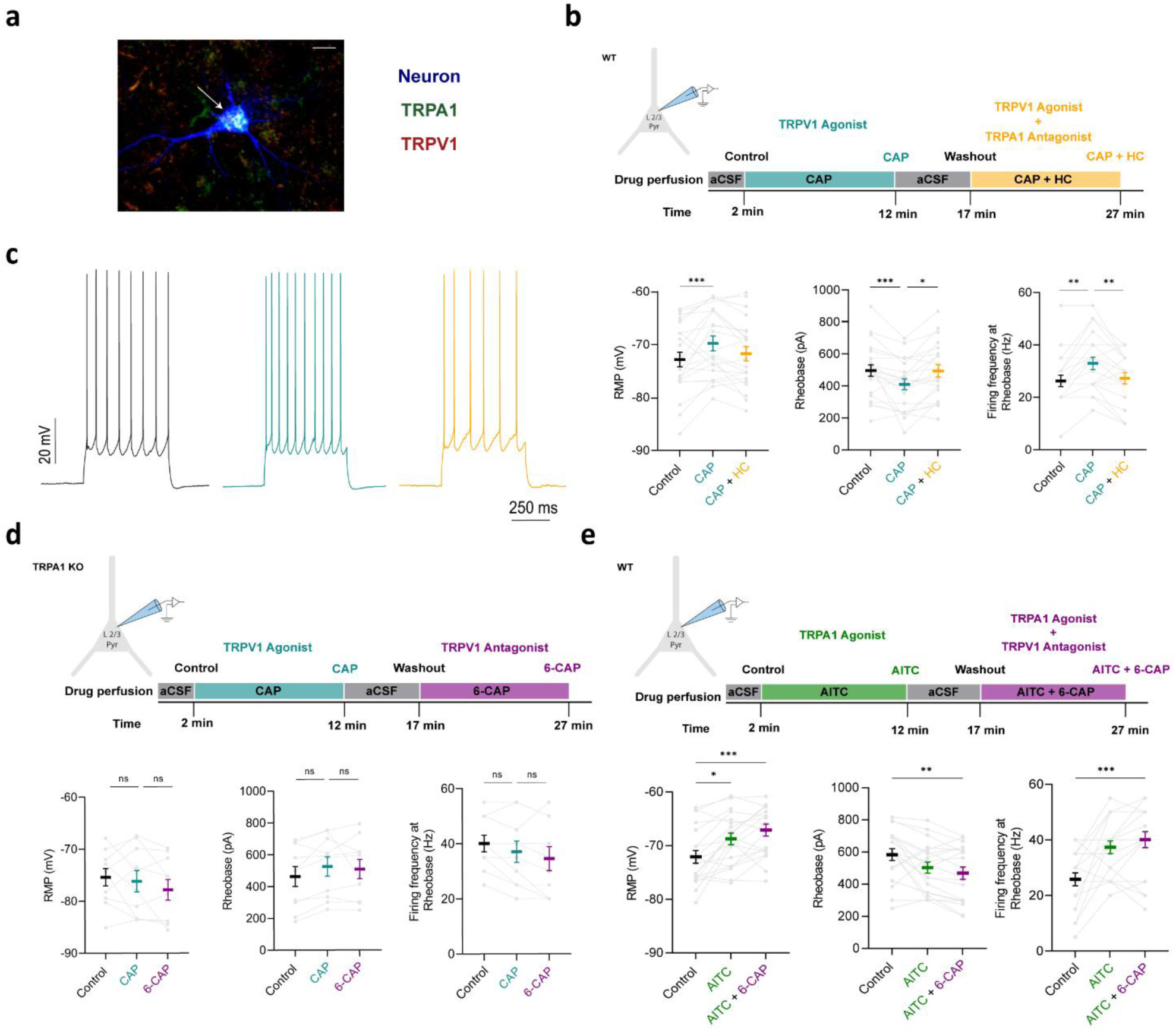
TRPV1-TRPA1 interactions modulate the excitability of cortical pyramidal neurons. **a)** Confocal images of a neurobiotin-filled cortical pyramidal neuron (blue) co-labelled with anti-TRPV1 (green) and anti-TRPA1 (red) antibodies, showing somatic co-expression of TRPV1 and TRPA1 on the recorded neuron (white arrow). Scale bar represents 10 μm. **b)** Top: Timeline of drug perfusions in wild-type (WT) mice. Bottom: Changes in resting membrane potential (RMP, left), rheobase (middle), and firing frequency at rheobase (right) following sequential application of aCSF, capsaicin (CAP; TRPV1 agonist), and a combination of CAP and HC-030031 (TRPA1 antagonist). Data from n = 22 neurons; (Friedman test with Dunn’s multiple comparisons). **c)** Example voltage traces from a neuron in response to a 400 pA current step under aCSF, CAP, and CAP + HC conditions. **d)** Top: Timeline of drug perfusions in TRPA1 knockout (KO) mice. Bottom: Changes in RMP (left), rheobase (middle), and firing frequency (right) across aCSF, CAP, and 6-CAP (TRPV1 antagonist) conditions. n = 10 neurons. Differences were not statistically significant (ns, Friedman test with Dunn’s multiple comparisons). **e)** Top: Timeline of drug perfusions in WT mice. Bottom: Changes in RMP (left), rheobase (middle), and firing frequency (right) upon application of aCSF, allyl isothiocyanate (AITC; TRPA1 agonist), and a combination of AITC and 6-CAP. n = 22 neurons; (Friedman test with Dunn’s multiple comparisons). Grey lines represent individual neurons; thick lines show the group mean. Error bars indicate SEM.ns =not significant, * p < 0.05, p **< 0.01, ***p < 0.001, ****p<0.0001

To assess functional coupling, we performed whole-cell current-clamp recordings in acute brain slices from wild-type mice. Activation of TRPV1 was sufficient to enhance intrinsic excitability *in vitro* (**Supplementary Figure 4**). To activate TRPV1, we applied the TRPV1 agonist capsaicin (CAP, 1 μM) for 10 minutes. This was then followed by co-application of CAP with a selective TRPA1 antagonist HC-030031 (HC, 10 μM; **Figure 3b**). CAP application produced a robust increase in neuronal excitability, evidenced by depolarisation of the resting membrane potential (RMP), a reduction in rheobase, and increased firing frequency at rheobase (**Figure 3b, c**). Critically, subsequent TRPA1 blockade significantly attenuated these effects. While CAP alone decreased rheobase and increased firing rate, co-application with HC reversed these changes, returning excitability measures toward baseline. These results indicate that TRPV1-dependent increases in excitability are contingent on intact TRPA1 function. To further validate the role of TRPA1, we repeated the CAP protocol in TRPA1 knockout mice. In contrast to the robust effects observed in wild-type neurons, CAP failed to elicit significant changes in RMP, rheobase, or firing frequency in TRPA1 KO neurons (**Figure 3d**). These findings provide convergent evidence that TRPA1 is necessary for TRPV1-mediated modulation of intrinsic excitability.

Next, we investigated whether the reverse dependency exists, specifically, whether TRPA1-driven excitability depends on TRPV1. Activation of TRPA1 using allyl isothiocyanate (AITC, 250 μM) produced a significant depolarisation of the RMP, decreased rheobase, and increased firing frequency (**Figure 3e**). Notably, these effects persisted in the presence of the TRPV1 antagonist 6-CAP and were preserved in TRPV1 knockout mice (**Supplementary Figure 5**). Together, these results reveal a directional and asymmetric functional interaction in which TRPV1-dependent modulation of excitability requires TRPA1, whereas TRPA1-mediated excitability occurs independently of TRPV1.

### TRPV1-TRPA1 interaction enhances cortical population coding and information transmission

Having established that TRPV1 modulates intrinsic excitability in a TRPA1-dependent manner in pyramidal neurons, we next examined whether this interaction influences sensory processing *in vivo*. To specifically characterise stimulus-evoked responses independently of behavioural variables, extracellular recordings were obtained from vS1 neurons in anaesthetised mice during application of mechanical whisker stimulation of varying amplitudes (0-160 *µ*m). Stimuli were delivered using a piezoelectric actuator, and cortical drug perfusion was applied sequentially via a lateral cannula targeting vS1 (**Figure 4a**). Neuronal responses were recorded under control conditions (aCSF), followed by application of the TRPV1 agonist (CAP), and subsequently CAP co-applied with the TRPA1 antagonist (CAP + HC).

**Figure 4.**
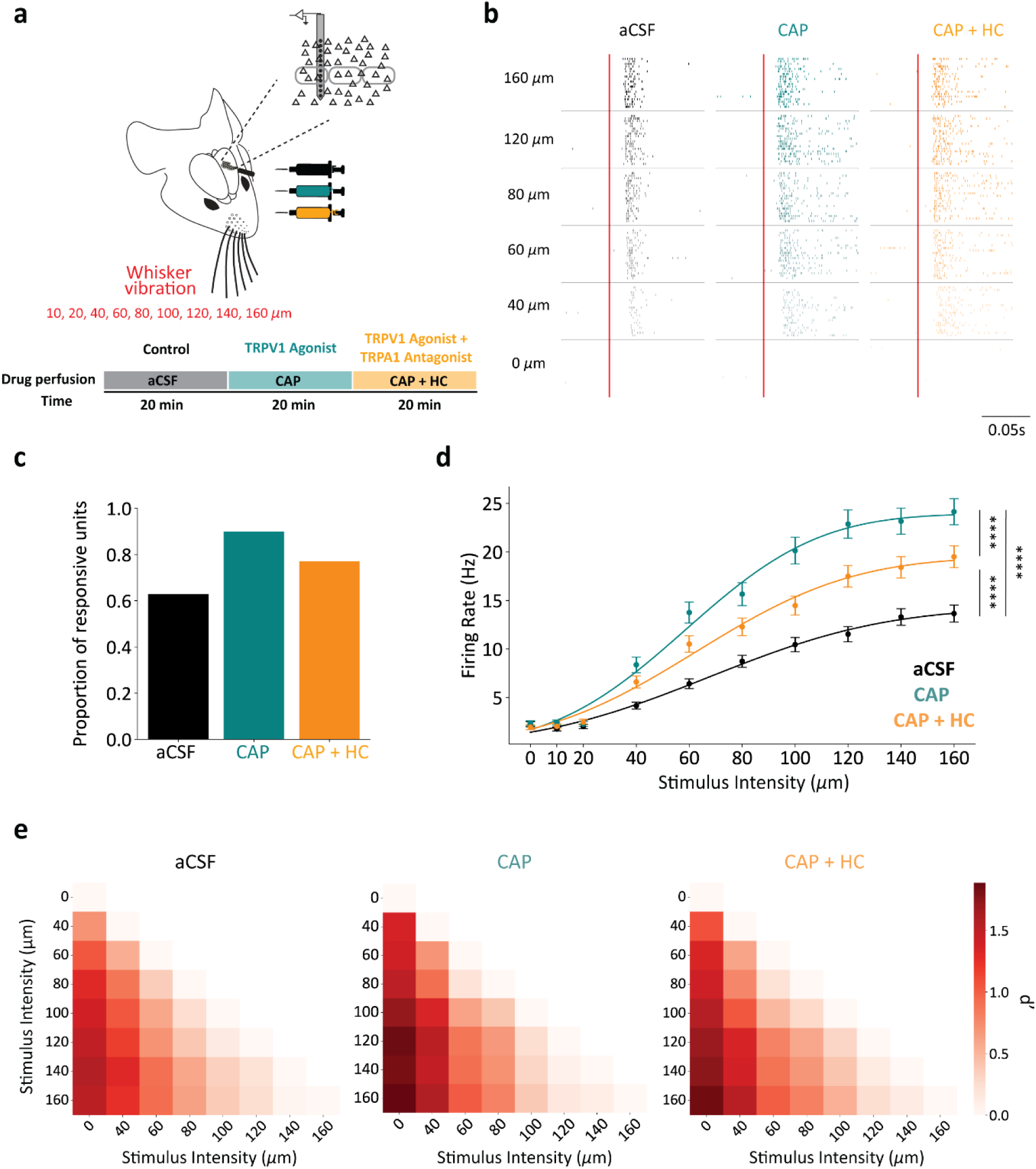
TRPV1-TRPA1 interaction modulates sensory representations in vS1. **a)** Schematic of the experimental design. Whisker stimulation was delivered using a piezoelectric actuator at varying intensities (0–160 μm). Cortical drug perfusion occurred in three sequential conditions: artificial cerebrospinal fluid (aCSF; control), capsaicin (CAP; TRPV1 agonist), and CAP co-applied with HC-030031 (TRPA1 antagonist). **b)** Raster plots of a representative vS1 neuron’s responses across the three conditions at multiple stimulus intensities. Each row represents a single trial; red vertical lines indicate stimulus onset. **c)** Proportion of neurons significantly modulated by whisker deflection under each condition. **d)** Population-averaged stimulus-evoked firing rate as a function of stimulus intensity across conditions (n = 358 neurons from 3 mice; ****p < 0.0001, RM one-way ANOVA with Tukey’s multiple comparisons). Data are presented as mean ± SEM. **e)** Heatmaps showing the average pairwise stimulus discriminability (d′) across all recorded neurons for aCSF (left), CAP (middle), and CAP + HC (right) conditions. Each matrix element represents the mean d′ for a given pairwise comparison of stimulus intensities, calculated per neuron and averaged across the population. Higher d′ values (darker red) indicate greater separability between response distributions.

Under all conditions, vS1 neurons exhibited graded responses to increasing stimulus intensities (**Figure 4b**). TRPV1 activation with CAP significantly increased the proportion of stimulus-responsive neurons relative to aCSF, an effect that was attenuated by introducing a TRPA1 inhibitor (**Figure 4c**). Similarly, population-averaged firing rates increased across the stimulus range following TRPV1 activation, reflecting a gain-like modulation of sensory responses. This enhancement was significantly reduced when TRPA1 was blocked (**Figure 4d**; p < 0.0001, RM one-way ANOVA with Tukey’s post hoc), indicating that TRPA1 contributes to TRPV1-mediated enhancement of sensory responsiveness.

To quantify changes in encoding precision, we computed the pairwise discrimination index (d′) across stimulus amplitudes for individual neurons (**Figure 4e**). Under control conditions, d′ values remained low for closely spaced amplitudes and increased primarily with larger amplitude separations. TRPV1 activation enhanced discriminability across a broader range of stimulus pairs, including those with small differences in amplitude. In contrast, co-application of CAP with the TRPA1 antagonist (HC) reduced these improvements at low stimulus amplitudes while sparing discriminability at higher amplitudes (**Figure 4e**). These results indicate that TRPA1 activity is required for TRPV1-mediated sharpening of sensory representations, particularly at low stimulus amplitudes.

The neuronal effects described above lead to two testable questions at the population level: (1) Does TRPV1 activation increase the signal-to-noise ratio, thereby enhancing stimulus representations particularly at low stimulus amplitudes? (2) Does TRPV1 alter the temporal precision and correlation structure within the neuronal population? Changes to population dynamics, such as reducing noise correlations or modulating the temporal envelope of responses, can strongly influence the fidelity of sensory encoding and improve downstream sensory readout ^20–24^. To address these questions, we examined how TRPV1 activation and its dependence on TRPA1 influence vS1 population coding by quantifying both the structure of shared variability and its impact on stimulus decoding accuracy.

Trial-to-trial covariability in neuronal activity (noise correlation) was estimated from pairwise spike count correlations under each condition ^25,26^. TRPV1 activation significantly altered the correlation structure within the neuronal population. Under CAP, positive noise correlations were significantly reduced relative to aCSF (p < 0.0001, Kruskal-Wallis test with Dunn’s multiple comparisons), while negative correlations were enhanced (p < 0.05, Kruskal-Wallis test with Dunn’s multiple comparisons; **Figure 5a**). Positive correlations typically impair population coding by reducing the effective dimensionality of neural responses and increasing redundancy, whereas negative correlations facilitate decorrelated activity and enhance stimulus separability ^20,21^. Blocking TRPA1 during TRPV1 activation (CAP + HC) reinstated positive correlations and diminished negative correlations, indicating that TRPA1 activity is required for TRPV1-dependent reorganisation of shared noise. These findings suggest that TRPV1-TRPA1 signalling enhances population-level discriminability by reducing shared variability and promoting beneficial decorrelation among neurons.

**Figure 5.**
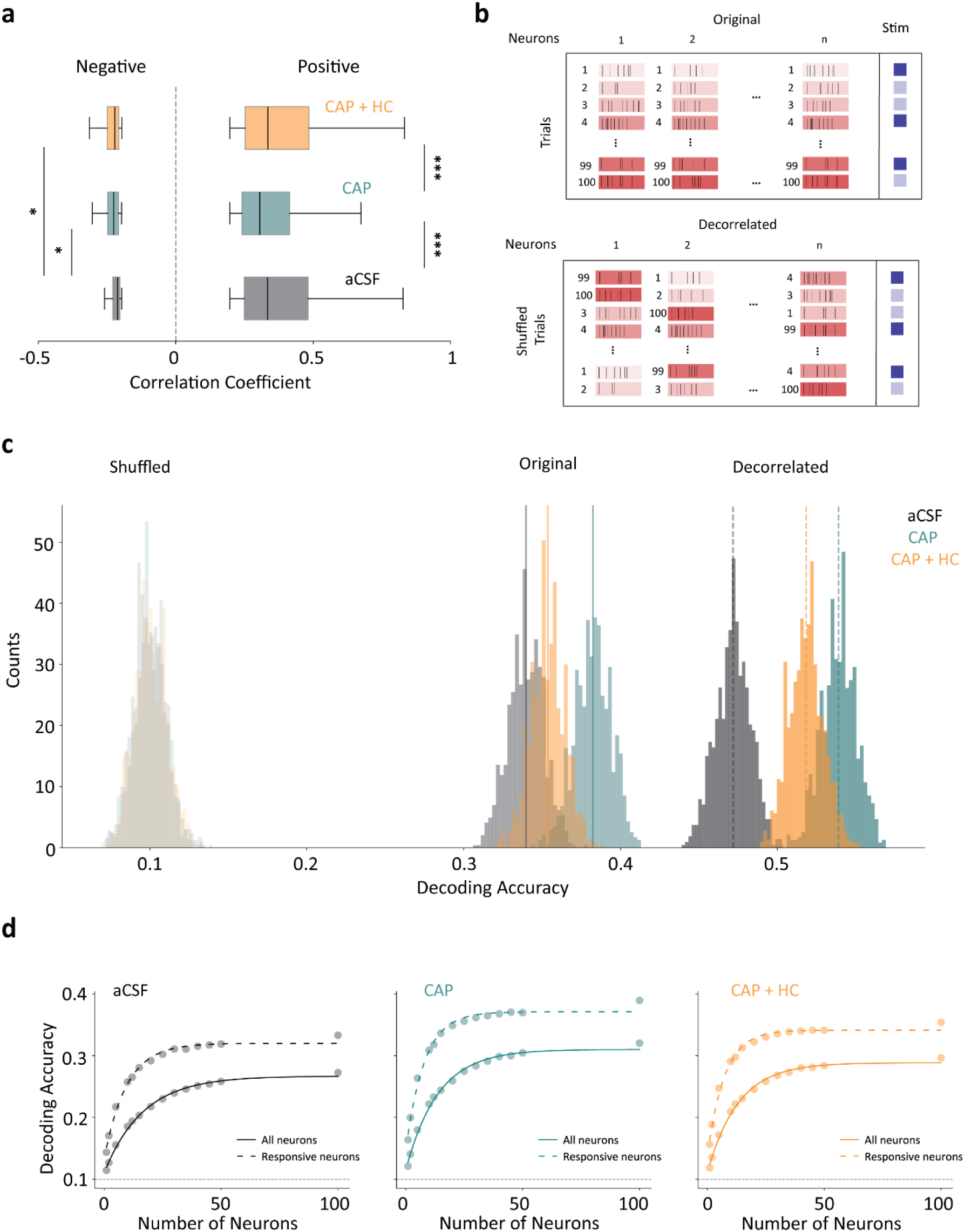
Population dynamics and the role of shared variability in stimulus encoding. **a)** Distribution of pairwise spike count correlation coefficients, separated into negative (left) and positive (right) correlations for each drug condition. Statistical comparisons were performed using the Kruskal-Wallis test with Dunn’s multiple comparisons; *p < 0.05, ***p < 0.0001. **b)** Schematic of spike raster plots based on simultaneously recorded neurons across trials in response to two different whisker stimuli (dark blue and light blue). Red boxes illustrate population activity across trials (rows) and neurons (columns). Left: Representation of original data preserving trial-to-trial covariability (noise correlations) between neurons. Right: Representation of decorrelated data generated by independently shuffling trial order for each neuron, preserving single-neuron response statistics while removing shared variability. **c)** Distribution of decoding accuracies across 5,000 iterations for shuffled (randomised control; left), original (middle), and decorrelated (right) trial structures under aCSF (grey), capsaicin (CAP; teal), and CAP + HC-030031 (orange) conditions for a representative mouse. Shuffled data reflect chance-level performance. Solid vertical lines indicate mean decoding accuracy for the original data; dashed lines indicate mean accuracy after trial shuffling. **d)** Decoding accuracy as a function of the number of neurons included in the population decoder for all three conditions. Solid lines show performance using pseudo-populations sampled from all recorded neurons; dotted lines show performance using only stimulus-responsive neurons.

To evaluate the functional impact of these changes in correlation, we trained a linear support vector machine (SVM) classifier to decode stimulus identity from trial-averaged firing rate vectors. This analysis quantifies how accurately population activity patterns distinguish between different stimuli across each condition. To isolate the contribution of noise correlation, we decorrelated the data by independently shuffling trial order for each neuron while preserving stimulus identity labels and firing rate distributions (**Figure 5b**). TRPV1 activation (CAP) significantly enhanced decoding accuracy compared to both aCSF and CAP + HC conditions (p < 0.0001, Kruskal-Wallis test with Dunn’s multiple comparisons; **Figure 5c, d; Supplementary Figure 7**). Removing shared variability through trial shuffling improved decoding accuracy across all conditions, demonstrating that noise correlation constrains information transmission. However, even after decorrelation, decoding performance remained higher under TRPV1 activation, indicating that TRPV1-TRPA1 signalling enhances sensory coding through both improved single-neuron selectivity and reorganisation of population structure.

We next examined decoding accuracy across pseudo-populations of varying sizes. Decoding accuracy under TRPV1 activation (CAP) reached asymptotic levels with fewer neurons than in other conditions (**Figure 5d**), indicating a more efficient distribution of sensory information across the population. Importantly, decoding accuracy under CAP remained higher even when restricted to stimulus-responsive neurons, suggesting that improved performance arises not from an increased number of active cells but from enhanced coding quality within individual neurons. Each neuron carries more distinctive, less redundant stimulus information, allowing the network to achieve the same level of accuracy with fewer contributing units. A similar trend was observed across the entire recorded neuronal population, although the overall decoding accuracy was lower due to larger variability (**Figure 5d**, solid lines). Together, these results demonstrate that TRPV1-TRPA1 interactions enhance cortical sensory representations by increasing excitability, sharpening stimulus discriminability and reorganising population dynamics to support more accurate and efficient sensory coding.

## Discussion

Across behavioural, imaging, electrophysiological and slice experiments, our findings converge on a model in which TRPV1 dynamically regulates cortical sensory processing through TRPA1-dependent increases in neuronal excitability. We demonstrate that activation of cortical TRPV1 enhances perceptual sensitivity, preferentially amplifies near-threshold sensory inputs and improves both single-neuron and population-level representations of tactile stimuli in awake mice. Our findings establish a direct link between TRPV1 signalling and sensory perception. More broadly, these findings challenge the traditional view of TRPV1 as a peripheral nociceptive receptor and instead identify it as a potential contributor to cortical sensory computation. The TRPV1-TRPA1 signalling axis therefore emerges as a novel mechanism by which endogenous ion channel signalling can dynamically regulate cortical gain and perceptual performance, extending the role of TRP channels beyond peripheral sensory transduction ^18,19,27^ to higher-order sensory processing.

Although TRPV1 is classically recognised as a peripheral detector of noxious heat, irritants and inflammation, numerous endogenous activators of the channel are present within the CNS. Endocannabinoids such as anandamide, oxidised lipids, reactive oxygen species and inflammatory mediators can regulate TRPV1 activity and are dynamically altered by physiological and pathological brain states ^9,28–30^. Our findings suggest that cortical TRPV1 functions not simply as a detector of external stimuli, but as an integrator of endogenous signalling pathways that modulate cortical sensitivity. Through this mechanism, changes in internal physiological state may influence sensory gain and the perceptual salience of external inputs.

The preferential enhancement of near-threshold tactile stimuli is consistent with cortical gain modulation, whereby neuronal responses are selectively amplified to optimise sensory representations without globally increasing network activity ^31–33^. Previously, we have shown that TRPA1 activation positively modulates sensory responses in both vS1 and V1 and enhances population-level sensory coding in the cortex ^34^. Our findings extend these observations by demonstrating that TRPV1-dependent modulation not only enhances sensory representations in the cortex but also directly affects perceptual sensitivity and behaviour in awake animals. Such gain modulation may underlie broad changes to sensory coding as induced by attention ^35–37^, expectation ^38–40^, or neuromodulation ^41–43^ and allow cortical circuits to dynamically adjust their sensitivity according to behavioural demands ^44,45^. In this context, TRPV1 signalling may provide a mechanism for dynamically tuning sensory gain in response to ongoing physiological states. Importantly, TRPV1 activation did not simply increase response magnitude but also improved sensory discriminability and cortical coding efficiency, suggesting that TRPV1-dependent modulation sharpens sensory representations rather than merely increasing cortical excitability.

Mechanistically, our data also support a model in which TRPV1 may engage TRPA1-dependent signalling pathways to regulate intracellular calcium dynamics and neuronal gain. In peripheral sensory neurons, TRPV1 and TRPA1 are extensively co-regulated through conserved intracellular signalling cascades involving phospholipase C (PLC), phosphatidylinositol 4,5-bisphosphate (PIP₂), protein kinase A (PKA), protein kinase C (PKC) and calcium-dependent signalling pathways ^46–51^. Activation of G-protein coupled receptors by inflammatory and neuromodulatory mediators can sensitise both channels through PIP₂ hydrolysis, phosphorylation-dependent mechanisms and increases in intracellular Ca²⁺, leading to enhanced neuronal excitability ^50–53^. We show that functional interactions between TRPV1 and TRPA1 extend to cortical neurons. and can regulate sensory coding. The requirement for TRPA1 in mediating TRPV1-induced increases in pyramidal neuron excitability further supports the existence of a conserved functional interaction between these channels within the cortex. Our findings are consistent with prior observations that cortical TRPV1 can adjust input-output transformations through multiple complementary mechanisms, including enhanced postsynaptic depolarisation ^17,54^, Ca²⁺ influx, modulation of presynaptic release probability ^55,56^, and regulation of inhibitory drive onto pyramidal neurons ^57–60^.

The intracellular signalling pathways that couple TRPV1 and TRPA1 also suggest a broader role for TRPV1 as a cortical neuromodulatory mechanism. Unlike classical neuromodulatory systems, such as cholinergic or noradrenergic pathways that originate from subcortical nuclei ^61–64^, TRPV1 is embedded directly within cortical neurons and can be locally recruited by endogenous ligands and intracellular signalling cascades. Consequently, TRPV1 signalling may provide a spatially restricted and state-dependent mechanism for regulating cortical gain. Fluctuations in arousal, metabolic state or neuroinflammatory signalling may directly influence sensory sensitivity by altering TRPV1-mediated modulation of cortical circuits.

At the population level, TRPV1 activation improved cortical coding efficiency by reducing shared variability between neurons. Reductions in noise correlation are known to enhance the information carried by neuronal populations by reducing shared variability and improving coding efficiency ^20,22,23,65,66^. In the rodent whisker system, active behavioural states are associated with reduced correlated neuronal activity and improved sensory representations ^67–71^. The ability of TRPV1 signalling to access similar coding regimes suggests that endogenous TRP channel activation may contribute to dynamic state-dependent regulation of cortical population activity. Further studies will be required to determine whether these effects arise through modulation of excitatory-inhibitory interactions or through recruitment of specific interneuron populations that regulate correlated variability within cortical networks ^72–74^.

Finally, our findings provide a conceptual link between the peripheral and central functions of TRP channels. In the peripheral nervous system, TRPV1 and TRPA1 act as polymodal sensors that detect environmental and internal signals and initiate sensory afferent activity ^75–77^. Here, we show that TRPV1 signalling persists at higher levels of the sensory hierarchy, shaping how sensory information is represented and perceived within the cortex. TRP channels may constitute a conserved molecular framework that operates across multiple stages of sensory processing, from detection of external stimuli in the periphery to modulation of cortical representations that determine perception. By bridging cellular excitability, population coding and behaviour, the TRPV1-TRPA1 signalling axis emerges as a plausible endogenous neuromodulatory mechanism that dynamically regulates cortical processing of sensory information and perceptual sensitivity. Whether TRPV1 exerts similar effects on information processing in other brain regions, or whether its central signalling contributes to additional physiological functions beyond sensory processing, remains to be determined.

## Methods

### Mice

Experiments were conducted using adult male and female C57BL/6J mice aged 28-42 weeks. Animals were group-housed (two mice per cage) in climate-controlled conditions with standard environmental enrichment and ad libitum access to food and a 12-hour reverse light/dark cycle in the behavioural suites at the John Curtin School of Medical Research (JCSMR). Mice undergoing behaviour were water-restricted, receiving water 2 hours after each session. All procedures were approved by the Animal Experimentation and Ethics Committee of the Australian National University (AEEC 2022/16).

### Surgery for 2P Ca²⁺ Imaging

Mice were anaesthetised with ∼2% isoflurane in oxygen, delivered through a nose mask at 0.--0.6 L/min. The depth of anaesthesia was monitored using respiratory rate and reflex tests. The eyes were protected using Viscotears gel. Mice were then placed in a stereotaxic frame, with body temperature maintained using a heating blanket. After excising the scalp, the skull was exposed, and periosteal tissue was gently removed using hydrogen peroxide to improve adhesion of dental cement.. The right vibrissal primary somatosensory cortex (vS1) was located at 3 mm lateral to the midline and 1.5 mm posterior to bregma. A 3 mm craniotomy was performed, followed by stereotaxic injection of the GCaMP7f Ca²⁺ indicator (Addgene, AAV1.Syn.GCaMP7f.WPRE.SV40). The injection was administered into the cortex at a depth of 230-250 μm from the dura across 4-6 sites, with four 32 nL injections per site, spaced 2-5 minutes apart and delivered at a rate of 92 nL/s. After the injections, a cranial window was covered with a 3 mm glass coverslip (0.1 mm thick, Warner Instruments, CT). A second, smaller craniotomy was performed lateral to vS1 to allow for the insertion of a guide cannula (26 Gauge, Protech International Inc.), which was secured in place using dental cement. Posterior to the cranial window, a head-bar was implanted using dental cement.

### Whisker stimulation

Vibration stimuli were delivered to the left (contralateral) whisker pad using a 2 cm × 2 cm mesh (aluminium) affixed to a ceramic piezoelectric actuator (Morgan Matroc, Bedford, OH). The mesh was positioned approximately 2 mm from the snout and oriented parallel to the whisker pad to ensure uniform engagement. Its position was kept constant across sessions by measuring the distance from the snout.

For calcium imaging experiments, stimuli consisted of 500 ms trains of Gaussian-shaped deflections, each lasting 15 ms with 10 ms intervals between deflections (40 Hz). Five stimulus amplitudes (0, 10, 20, 40, and 80 µm) were presented in pseudorandomised order, with interstimulus intervals (ISI) of 5-10 seconds. In electrophysiology experiments, individual 20 ms Gaussian-shaped deflections were delivered at ten discrete amplitudes (0, 10, 20, 40, 60, 80, 100, 120, 140, and 160 *µ*m) with ISIs of 0.4-1 second.

All stimulus waveforms were generated and controlled using MATLAB scripts and delivered via a National Instruments data acquisition board (PCIe-6321 for electrophysiology) at a 20 kHz sampling rate. Variability due to non-immobilised whiskers was consistent across conditions, ensuring valid relative comparisons.

### Drug perfusion

Drugs were delivered through the implanted guide cannula via a syringe pump (CMA402, Harvard Apparatus, USA) at a rate of 0.7 μL/min during the session. For pharmacological manipulation, in a given session, either artificial cerebrospinal fluid (aCSF) or 1 µM capsaicin (CAP; TRPV1 agonist) or 10 µM 6’nordihydroiodocapsaicin (6-CAP; TRPV1 antagonist) was perfused. The concentration of the TRPV1 agonist used was based on previously published dose-response curves and has been reported to be optimal for TRPV1 activation (Hazan et al., 2015). The order of drugs was pseudo-randomised across sessions.

### Habituation and Behavioural Training

To enable controlled whisker stimulation and behavioural monitoring, mice were head-fixed in a custom-designed apparatus. Animals were placed in a clear acrylic tube (4 cm inner diameter), with their heads protruding from the front to allow forepaw support on the tube’s edge. Using a surgically implanted custom head bar, the animal was secured in a stable yet natural crouched position, allowing unobstructed whisker access to the stimulus apparatus. Licking behaviour and reward delivery were monitored using a steel lick-port positioned approximately 0.5 mm below the lower lip and ∼5 mm posterior to the nose tip. Licks were detected using a capacitive sensor via an Arduino Uno microcontroller (Duinotech Classic, Cat#XC4410), which transmitted voltage signals to a National Instruments data acquisition system. Detection thresholds were software-defined, and each detected lick triggered a gravity-fed solenoid valve to dispense a drop of 5% sucrose solution. All behavioural control, stimulus delivery and data acquisition were implemented with MATLAB scripts. Signals from the lick-port and stimulus hardware were acquired at 100 kHz via a National Instruments DAQ board, ensuring precise synchronisation and event timing.

After a 7-day post-surgery recovery period, mice were introduced to water restriction, during which they habituated to the experimenter and the head-fixation setup. Habituation began with brief periods in the acrylic tube, which were progressively increased in duration. Once accustomed, animals were gently restrained near the headpost using haemostatic forceps, with hold times extended over sessions. During this period, mice received small drops of 5% sucrose as positive reinforcement and were given 2 hours of unrestricted water access after each session. Training for the detection task began once mice tolerated full head fixation. In the initial phase, each lick on the reward spout triggered delivery of a sucrose drop, reinforcing the lick-reward association. In the second phase, a vibration stimulus was continuously presented until the animal produced three licks, after which a reward was dispensed. This was followed by a 60-second no-go period with no stimulus or reward, encouraging temporal association between stimulus and reward availability. Next, the go/no-go discrimination task was introduced. In each trial, mice were presented with either a go stimulus (80 µm vibration) or a no-go condition (0 µm), with a variable inter-trial interval (5-10 s) to reduce anticipatory behaviour. Upon achieving high performance (>80% correct responses - Hit and CR), the vibration duration was reduced from 1s to 500 ms.

Finally, animals were transitioned to a detection task using five vibration amplitudes (0, 10, 20, 40 and 80 µm), presented in pseudo-randomised blocks of five trials (one per amplitude). Each imaging session comprised 100 trials, with only one session conducted per day to prevent fatigue or adaptation. This task was repeated across three sessions for each drug condition (aCSF, CAP and 6-CAP), allowing for within-subject comparisons across pharmacological conditions. During the task, neuronal activity was also recorded using a 2-photon microscope.

### 2P Ca²⁺ Imaging

Three to four weeks after intracortical injection of GCaMP7f, Ca^2+^ imaging was performed using a 2-photon microscope system (ThorLabs, MA), equipped with a Ti:Sapphire laser (Chameleon, Coherent) tuned to 920 nm. Imaging was conducted through a 16× water-immersion objective lens (0.8 NA, Nikon), with the focal plane positioned in cortical layers 2/3. Neuronal activity was recorded as Ca²⁺ transients at a resolution of 512 × 512 pixels and a frame rate of approximately 30 Hz. Laser power was adjusted between 40 and 75 mW depending on the level of GCaMP7f expression in each animal. Images were acquired using ThorImage software (ThorLabs), and all frames were time-locked to stimulus presentation via a National Instruments data acquisition system. Neurons were tracked across sessions, allowing recordings to be performed from the same neurons under the three different drug conditions.

### Behavioural Analysis

Trials were classified as Hits if the animal performed at least one lick within the 0-500 ms window following stimulus onset, and no licks were detected in the 500 ms preceding the stimulus. Miss trials were defined by the absence of licking in the 0-500 ms following stimulus onset. Lick frequency was measured by subtracting the number of licks in the 500 ms before stimulus presentation from the number of licks in the 1-second period after stimulus, giving an indicator of stimulus-evoked licking behaviour. Hit and False Alarm (FA) rates were first converted to z-scores by applying the inverse normal transformation, which maps response probabilities onto a standard normal distribution. D-prime (d′) was then calculated as the difference between the z-scored Hit rate and the z-scored FA rate as:

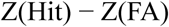

To link trial-to-trial variability in cortical activity to behavioural outcomes, we employed a stratified (stimulus-balanced) resampling approach. Trials were pooled across animals and repeatedly subsampled (100 trials per iteration; 25 per stimulus intensity at 10, 20, 40, and 80 µm). For each subset, behavioural performance was quantified as the stimulus-balanced hit rate, and neuronal activity as the mean normalised Ca^2+^ response (AUC). Repeating this procedure 1,000 times yielded paired behavioural and neural measures, and their association was assessed using Pearson’s correlation.

### Neuronal Analysis (2-Photon Ca²⁺ Imaging)

2-photon imaging data were processed using the Suite2P Python package (https://github.com/cortex-lab/Suite2P) for motion correction and semi-automated detection of regions of interest (ROIs). Identified ROIs were manually curated using ImageJ to ensure anatomical accuracy. Neuropil signals were subtracted from each neuron’s Ca^2+^ trace using a custom Python script to minimise background contamination.

Fluorescence changes were quantified as ΔF/F₀, where F₀ represented the baseline fluorescence, calculated as the average fluorescence over the 500 ms preceding stimulus onset. Neurons were classified as responsive if their stimulus-evoked ΔF/F₀, measured during the 500 ms following stimulus onset, was significantly greater than baseline activity (Wilcoxon signed-rank test, p < 0.05). For each responsive neuron, the area under the curve (AUC) of the ΔF/F₀ trace was calculated over a 1-second window from stimulus onset. AUC was chosen over peak ΔF/F₀ because it captures both the magnitude and temporal extent of stimulus-evoked calcium responses, and the integration window was selected to cover the entire post-stimulus response period. In our dataset, relatively few neurons exhibited decreases in ΔF/F₀, and their number was insufficient to permit robust statistical analysis. Accordingly, subsequent analyses focused on stimulus-evoked increases in population activity.

Due to the strong dependence of detection probability on stimulus amplitude, behavioural outcomes were not balanced across stimulus levels. Analyses were therefore designed to minimise sensitivity to trial count imbalances.

To isolate stimulus responses, “*stimulus index*” was calculated using AUC from Miss and Correct Rejection (CR) trials as:

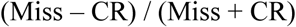

To isolate the choice element, “*choice index*” was calculated using AUC from FA and CR trials as:

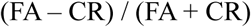

Neurons were classified as significantly stimulus- or choice-modulated if their observed index exceeded the 95% confidence interval of the shuffled null distribution, based on a two-sided permutation test with p < 0.05. Neurons were tracked across imaging sessions to evaluate consistency in stimulus and choice encoding under different experimental conditions.

To evaluate the discriminability of sensory representations, a “Discrimination Index” was calculated as follows:

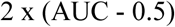

where the AUC was obtained from receiver operating characteristic (ROC) curves comparing neural activity between stimulus-present and stimulus-absent trials at each vibration intensity, using only Miss and Correct Rejection trials to minimise licking and movement-related confounds in the calcium signals.

### Surgery for extracellular electrophysiology recording

Mice were initially anaesthetised with isoflurane for induction and subsequently maintained under anaesthesia via intraperitoneal injection of urethane (1.1 g/kg) combined with chlorprothixene (5 mg/kg). The depth of anaesthesia was routinely monitored by assessing hind paw withdrawal and corneal reflexes and maintained at a stable level with supplemental doses of urethane (10% of the initial dose) administered as required.

Following induction, the animal was secured in a stereotaxic frame using ear bars and placed on a feedback-controlled heating blanket to maintain body temperature. After excising the scalp, the skull was exposed, and periosteal tissue was gently removed using hydrogen peroxide to improve adhesion of dental cement. The right vibrissal primary somatosensory cortex (vS1) was located using stereotaxic coordinates (3 mm lateral to the midline, 1.5 mm posterior to bregma) and a 3 mm craniotomy was performed over this area. A second, smaller craniotomy was made lateral to vS1 to allow insertion of a guide cannula, which was fixed in place using dental cement. Posterior to the craniotomies, a head-bar was implanted using dental cement.

### Extracellular electrophysiology recording

Following surgery, mice were transferred to the recording setup and head-fixed. Body temperature was maintained using a heating pad throughout the experiment. Extracellular recordings were performed using a high-density Neuropixels probe (1.0), which contains 960 electrode sites along a 1 cm shank, with 384 channels active simultaneously for recording. Signals were bandpass filtered (250–5000 Hz) and digitised at a 30 kHz sampling rate, then continuously recorded to a disk. The probe was carefully lowered into the brain at a 45° angle using a micromanipulator and allowed to settle for approximately 20 minutes prior to the start of data acquisition.

### Spike sorting and population analyses

Data was sorted offline using the Kilosort package (https://github.com/MouseLand/Kilosort), an automated clustering algorithm that detects and classifies spikes based on waveform similarity. The resulting clusters were manually curated using Phy GUI (https://github.com/cortex-lab/phy). To identify individual units, spike clusters were analysed by their waveform shapes, cross-correlograms and spatial distribution across recording channels. Clusters were manually refined by grouping or splitting them as required. The final classification as individual units was confirmed from consistent waveform patterns and clear refractory periods in auto-correlograms. Following spike sorting and unit identification, neuronal responses were quantified using peristimulus time histograms (PSTHs) constructed using 5 ms bins. To identify responsive neurons, we examined the activity within a 50 ms post-stimulus window. If 2 consecutive bins within this window exceeded the mean pre-stimulus activity (calculated over 50 ms before stimulus onset) by more than 2 SDs, the neuron was classified as responsive to that stimulus intensity under that condition. For population analyses, we included neurons responsive to any stimulus under any condition, as indicated by the n values on the figures. The mean firing rate was calculated using spikes occurring within the 500 ms post-stimulus window. The peak firing rate was defined as the maximum firing rate observed in any single 5 ms bin within the same post-stimulus window. Noise correlations were quantified using Pearson correlation coefficient.

### d’

To quantify stimulus discriminability at the single-neuron level, we computed d-prime (d′), a signal detection theory metric that incorporates both the difference in mean responses and the trial-to-trial variability. d′ values were calculated for all pairwise combinations of stimulus intensities using net evoked firing rates, defined as the average firing rate within a fixed post-stimulus window. For each stimulus pair, d′ was computed per neuron as:

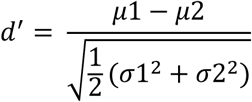

where μ1 and μ2 are the mean firing rates for the two stimuli, and σ_1_ ^2^ and σ ^2^ are the variances in firing rates across trials for each stimulus. This analysis quantifies the separability of neuronal response distributions across stimulus conditions while accounting for variability in evoked activity.

### Decoding accuracy using SVM

To evaluate the extent to which stimulus intensity could be predicted from evoked spike activity under each drug condition, neural decoding was performed using a linear support vector machine (SVM) classifier similar to Koren 2021 ^78^. For each condition and stimulus intensity, spike counts were recorded within a 100 ms post-stimulus window across all responsive units, resulting in a high-dimensional feature matrix composed of spike counts per trial. The decoding target was stimulus intensity, and features consisted of spike counts from all responsive neurons for each trial. Decoding was conducted separately for each drug condition using 5,000 iterations of classification. In each iteration, data were randomly partitioned into training (80%) and testing (20%) sets using stratified 5-fold cross-validation to preserve the distribution of stimulus classes. The decoding accuracy for each iteration was computed as the mean classification accuracy across the five folds and the final accuracy for each condition was determined as the average accuracy across all iterations (Koren, 2021). To remove noise correlation, spike count vectors were decorrelated by shuffling within trials for the same stimulus and drug condition prior to decoding, and accuracy was recalculated using the same procedure. A shuffle control was performed by randomly permuting stimulus labels across all trials, thereby establishing a chance-level decoding accuracy. Here, “shuffled” refers to a random permutation of stimulus labels across trials, which removes the relationship between neural activity and stimulus identity, thereby abolishing stimulus-specific information. In contrast, “decorrelation” refers to shuffling neuronal responses across trials within the same stimulus and condition, preserving stimulus labels while selectively removing correlated variability among neurons.

### Statistical analysis

Statistical analyses were performed using appropriate tests based on data distribution. Normality was assessed using the Kolmogorov–Smirnov test. For repeated-measures comparisons, an RM one-way ANOVA with Tukey’s multiple comparisons was used when data were normally distributed, otherwise Friedman or Kruskal-Wallis (independent) test followed by Dunn’s post hoc test was applied. Results are presented as mean ± standard error of the mean (SEM), unless otherwise specified.

### Whole-cell electrophysiology and immunohistochemistry

Mice were anesthetised with isoflurane and rapidly decapitated. Brains were extracted and placed in ice-cold, carbogen-bubbled slicing solution (110 mM choline chloride, 11.6 mM Na–ascorbate, 7 mM MgCl₂, 0.5 mM CaCl₂, 3.1 mM Na-pyruvate, 2.5 mM KCl, 1.25 mM NaH₂PO₄, 26 mM NaHCO₃, 10 mM glucose, pH 7.4). Coronal cortical slices (300 µm) were prepared using a Leica Vibratome and incubated in carbogen-enriched solution (92 mM NaCl, 2.5 mM KCl, 1.2 mM NaH₂PO₄, 2 mM CaCl₂, 2 mM MgSO₄, 25 mM glucose, 3 mM Na-pyruvate, 30 mM NaHCO₃, 20 mM HEPES, osmolarity ∼300–310 mOsm) at 34 °C for 30 minutes, then at room temperature for 30 minutes.

Recording pipettes were pulled from borosilicate glass (1.5 mm OD, 0.86 mm ID, 5–7 MΩ) and filled with internal solution (130 mM K-gluconate, 10 mM KCl, 10 mM HEPES, 4 mM MgATP, 0.3 mM Na₂GTP, 10 mM Na₂-phosphocreatine, 0.2% neurobiotin, pH 7.25, osmolarity ∼280 mOsm). Slices were perfused with carbogen-bubbled aCSF (125 mM NaCl, 3 mM KCl, 1.25 mM NaH₂PO₄, 25 mM NaHCO₃, 25 mM glucose, 2 mM CaCl₂, 1 mM MgCl₂) at 2–3 mL/min and 30–34 °C. Layer 2/3 and 5 pyramidal neurons were visualised under DIC microscopy. Whole-cell recordings were established in voltage clamp at –70 mV and switched to current clamp; series resistance was 15-22 MΩ. Firing properties were assessed using current steps (–300 to 500 pA, 750 ms) and ramp currents (–200 to 700 pA, 1500 ms). TRPA1 (AITC 250 µM, HC-030031 10 µM) and TRPV1 (capsaicin 1 µM, 6’-CAP 10 µM) modulators were applied for 10 minutes before testing. Signals were filtered at 10 kHz and digitised at 50 kHz using Axograph.

After recording, slices were fixed in 4% paraformaldehyde and processed for immunohistochemistry. Sections were permeabilized in 1% Triton X-100, blocked with 2% BSA, and incubated with primary antibodies against TRPA1 (rabbit, 1:200, 45 h) and TRPV1 (guinea pig, 1:200, 45 h), followed by fluorescent secondary antibodies (donkey anti-rabbit Alexa 555, 1:1000; goat anti-guinea pig Alexa 647, 1:500). Neurobiotin-filled neurons were visualized with streptavidin-AlexaFluor 488 (3 h, 4 °C). Sections were mounted and imaged on a Nikon A1 confocal microscope (1 *µ*m Z-stacks, excitation 488, 555, 647, 405 nm).

### Data Analysis

Recordings from healthy, stable neurons were analysed in Axograph. Resting membrane potential and rheobase were measured from ramp currents, and firing frequency and action potential properties (height, 20-80% rise time, half-width) were extracted from current step responses. All statistical analyses were performed using GraphPad Prism 9. Data were assessed for normality using the Shapiro-Wilk test. For repeated-measures comparisons, an RM one-way ANOVA with Tukey’s multiple comparisons was used when data were normally distributed, otherwise Friedman test followed by Dunn’s post hoc test was applied. Results are presented as mean ± standard error of the mean (SEM), unless otherwise specified.

## Acknowledgments

This work was supported by the Australian Research Council (ARC) Discovery Project Grant (DP240103043) and the ARC Centre of Excellence for Integrative Brain Function (CE140100007) awarded to E.A., and by the NHMRC Investigator Grant (GNT2016513) and Ideas Grant (GNT1181643) awarded to E.K.

## Author Contributions

L.A., E.A., and E.K. conceived and designed the project. L.A. performed all experiments, with S.G. assisting with Neuropixels recordings. L.A. analysed all data, and L.A. and K.S. analysed the Neuropixels data. L.A. wrote the manuscript, and all authors edited and approved the final version of the manuscript.

## Data availability

The data that support the findings of this study are available from the corresponding authors upon reasonable request.

## Code availability

All custom-written python and MATLAB scripts used for data acquisition and analysis will be made available by the authors upon request.

