## Supplementary Figures for "Cortical TRPV1 modulates sensory processing through TRPA1-dependent mechanisms"

#### Supplementary Figure 1

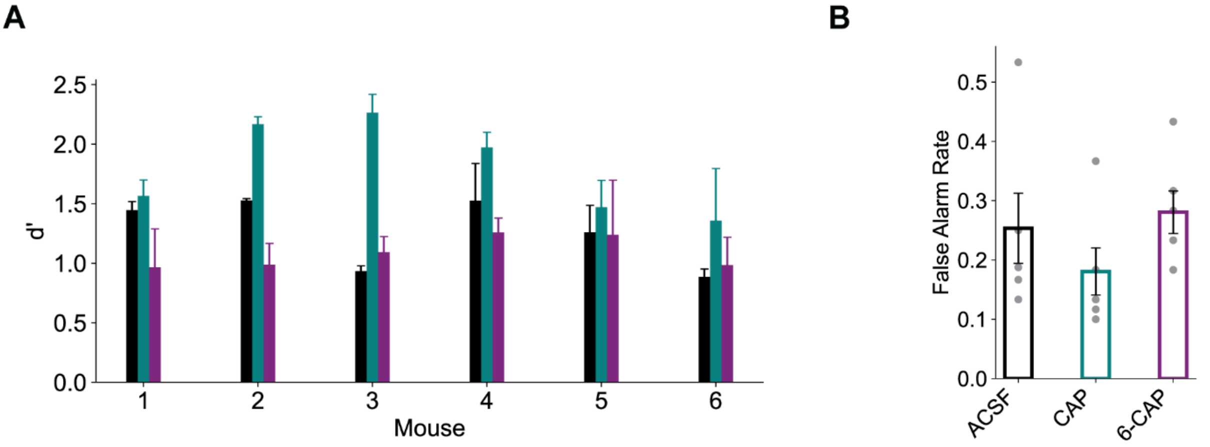


***Supplementary Figure 1. TRPV1 modulation of sensory perception (A)*** *Average d-prime (d') values across all stimuli for each mouse and under ACSF (black), CAP (teal) and 6-CAP (purple).* ***(B)*** *False Alarm rates across all mice under all conditions. Bars represent mean and error bars represent SEM.*

#### Supplementary Figure 2

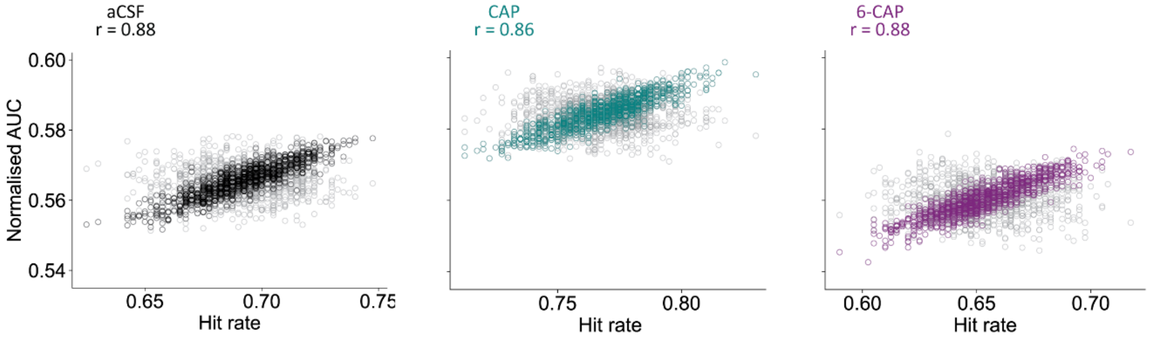


***Supplementary Figure 2. TRPV1 modulation of sensory perception*** *Shared variability between behavioural performance and vS1 activity assessed by stimulus-balanced resampling. Each point represents a resampled subset of 100 trials, plotted as hit rate versus mean normalised* $Ca^{2+}$ *response (AUC). Coloured circles show real resampled data and grey circles indicate shuffled controls. Pearson correlation coefficients (r) are shown.*

#### Supplementary Figure 3

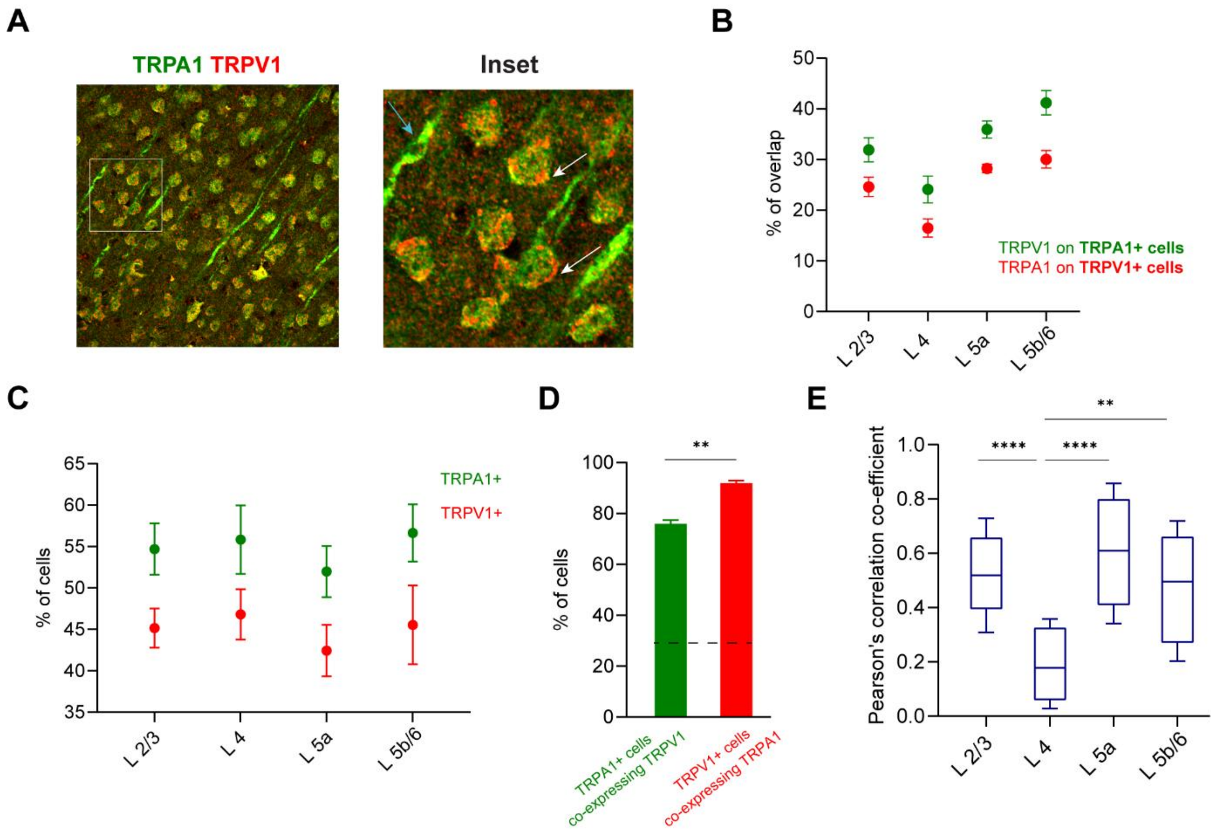


***Supplementary Figure 2. TRPV1-positive cells highly co-express TRPA1 (A)*** *A cortical section representing overlap (yellow) between TRPA1 (green) and TRPV1 (red) (left inset). Right inset represents the magnified view of cells from the white boxed area in the merged image (left inset), indicating the co-expression of TRPA1 and TRPV1 in the soma (white arrow) and lack of that co-expression on the dendrite (blue arrow) (right inset). Scale bars are 50 µm* ***(B)*** *Mean percentage of overlap between TRPA1 and TRPV1 across individual cortical layers, measured as Manders' coefficient M1 (fraction of TRPA1 overlapping TRPV1; $p<0.0001$; One-way ANOVA) and M2 (fraction of TRPV1 overlapping TRPA1; $p<0.0005$; One-way ANOVA) from confocal images (n=15)* ***(C)*** *Mean percentage of cortical cells expressing TRPA1 ($p=0.0110$; One-way ANOVA) and TRPV1 ($p= 0.0291$; One-way ANOVA) across all layers (n=15)* ***(D)*** *Mean percentage of TRPA1-positive cells co-expressing TRPV1 (green) and TRPV1-positive cells co-expressing TRPA1 (red) (n=15; $p>0.05$; Two-way ANOVA). Dotted line represents probability of co-expression* ***(E)*** *Box and whisker plot representing the Pearson's correlation coefficient across individual cortical layers (L2/3, L4, L5a and L5b/6). Box indicates the first to third quartile range with the median represented by the line in the middle. The whiskers represent minimum and maximum Pearson's correlation value (n=15; $p<0.0001$; One-way ANOVA). Error bars represent SEM. * $p < 0.05$, ** $p < 0.01$, *** $p < 0.001$*

#### Supplementary Figure 4

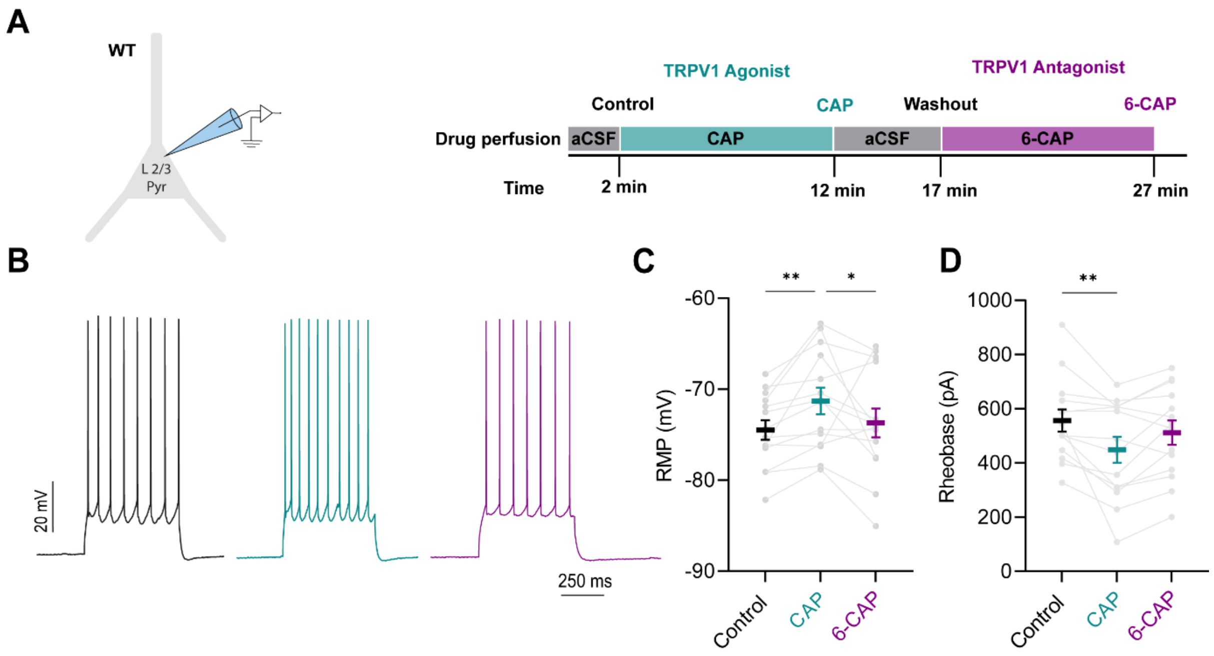


***Supplementary Figure 3. TRPV1 activation enhances the excitability of cortical layer 2/3 pyramidal neurons.*** ***(A)*** *Schematic of experimental protocol showing sequential drug perfusions during somatic whole-cell recordings from cortical layer 2/3 pyramidal neurons in wild-type (WT) animals. Recordings began 2 minutes after achieving whole-cell configuration (control), followed by a 10-minute application of 1 µM CAP (TRPV1 agonist), a 5-minute washout with aCSF and a 10-minute application of 10 µM 6-CAP (TRPV1 antagonist).* ***(B)*** *Example voltage responses to a 300 pA current step under control (black), CAP (blue) and 6-CAP (purple) conditions.* ***(C)*** *Resting membrane potential (RMP; n=14; p=0.0035) and* ***(D)*** *rheobase (n = 14, p = 0.0464) across the three conditions. Grey lines represent individual neurons, coloured lines represent mean. Error bars denote SEM. Friedman test with Dunn's multiple comparison test. *p < 0.05, **p< 0.01.*

### Supplementary Figure 5

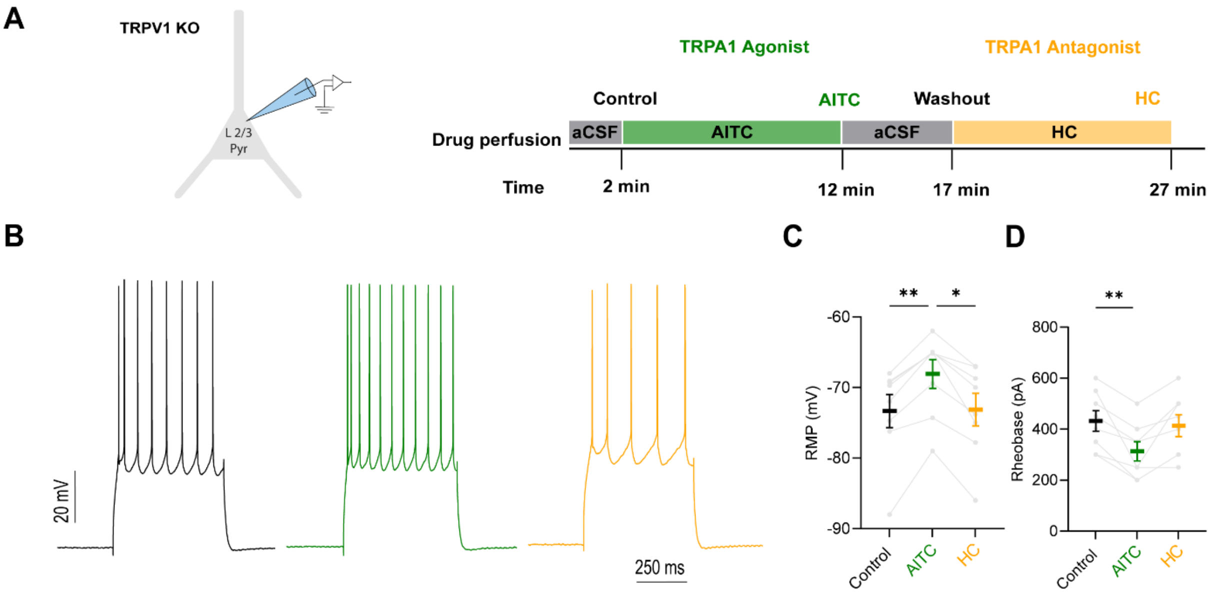


***Supplementary Figure 5. TRPA1 modulation is non-reliant on a functional TRPV1. (A)*** *Schematic of experimental protocol showing sequential drug perfusions during somatic whole-cell recordings from cortical layer 2/3 pyramidal neurons in TRPV1 Knockout (KO) animals. Recordings began 2 minutes after achieving whole-cell configuration (control), followed by a 10-minute application of 250 µM AITC (TRPA1 agonist), a 5-minute washout with aCSF, and a 10-minute application of 10 µM HC (TRPA1 antagonist).* ***(B)*** *Example voltage responses to a 500 pA current step under control (black), AITC (green) and HC (orange) conditions.* ***(C)*** *Resting membrane potential (RMP; n=8; p=0.0009) and* ***(D)*** *rheobase (n = 8; p= 0.0003) across the three conditions. Grey lines represent individual neurons, coloured lines represent mean. Error bars denote SEM. Friedman test with Dunn's multiple comparison. ns = not significant*

#### Supplementary Figure 6

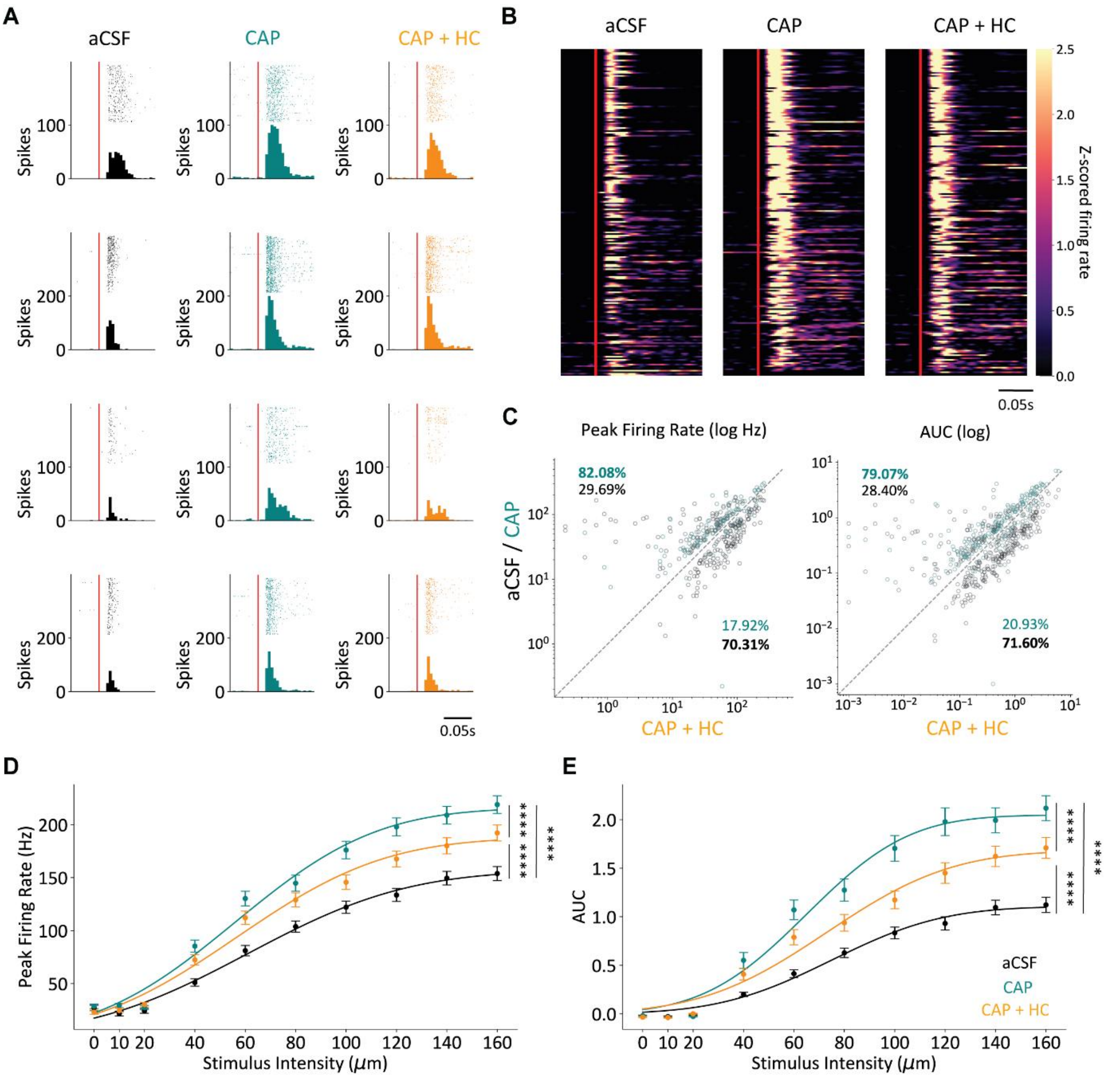


***Supplementary Figure 6. Temporal dynamics of evoked sensory responses across pharmacological conditions. (A)*** *Peri-stimulus time histograms (PSTHs, 5-ms bins) and single-trial raster plots from representative neurons under aCSF (black), CAP (teal), and CAP + HC (orange) in response to 100 μm whisker stimulation (stimulus onset: red line).* ***(B)*** *Heatmaps of z-scored firing rates across all recorded neurons, sorted by peak response under aCSF.* ***(C)*** *Scatter plots comparing peak firing rate (left) and response area under the curve (AUC, right) for individual units under aCSF vs CAP + HC (black) and CAP vs CAP + HC (teal). Percentages denote the proportion of neurons in each quadrant.* ***(D)*** *Population-averaged peak firing rate as a function of stimulus intensity under each condition (n=358 neurons from 3 mice, $p<0.0001$ between conditions)* ***(E)*** *Population-averaged AUC across stimulus*

*intensities (n=358 neurons from 3 mice, p<0.0001 between conditions). Data represent mean ± SEM. Statistical comparisons performed using repeated-measures one-way ANOVA with Tukey's multiple comparisons; ****p <0.0001.*

#### Supplementary Figure 7

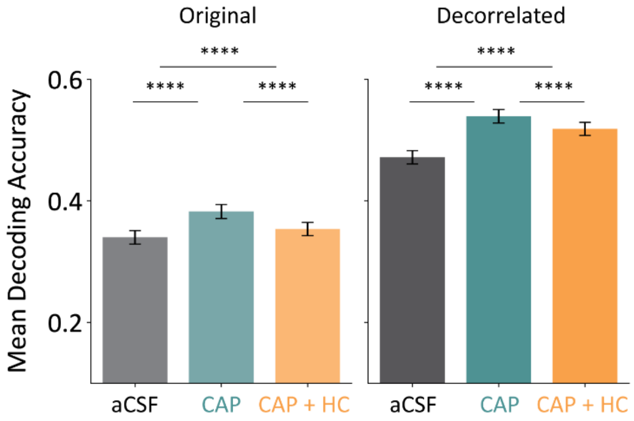


***Supplementary Figure 7. Noise correlations constrain stimulus encoding.*** *Mean decoding accuracy under each condition for original (left) and decorrelated (right) data. Data represent mean ± SEM. Statistical comparisons were performed using Kruskal–Wallis test with Dunn's multiple comparisons; ****p<0.0001.*
